# Characterizing trajectory-like chromatin architectures with Fun2

**DOI:** 10.1101/2025.02.25.640072

**Authors:** Zhengrong Zhangding, Xuhao Liu, Jiazhi Hu

## Abstract

Chromatin is intricately folded into dynamic 3D structures, orchestrating key DNA metabolic processes. DNA replication, a core chromatin metabolic event, is tightly linked to these chromatin architectures. Recently, a replication-associated chromatin interaction structure was discovered as fountains via Replication-associated *in situ* Hi-C (Repli-HiC), supporting that replication forks remain spatially coupled from initiation to termination. Chromatin fountains are pivotal for understanding DNA replication within the complex chromatin landscape. However, the characteristics of these fountains can vary due to factors such as origin firing efficiency and local chromatin organization. In this study, we introduce a reinforcement learning-based computational framework, integrated with Monte Carlo Tree Search (MCTS) and value gradient optimization, to develop the Fun2 algorithm. Fun2 enables a comprehensive characterization of trajectory-like chromatin architectures, including both chromatin fountains and stripes, with enhanced adaptability and accuracy. This tool facilitates systematic investigation of the spatiotemporal dynamics of DNA replication and extends to other chromatin remodeling processes, such as Cohesin-mediated loop extrusion. The Fun2 algorithm provides a versatile computational tool for deciphering dynamic chromatin architectures, offering new insights into genome organization and its regulatory mechanisms.

## Introduction

The 3D architectures of chromatin serve as the structural framework for a variety of essential DNA metabolic processes, including mediating enhancer-promoter interactions in transcription regulation^1–5^ and the scanning process of recombination-activating gene (RAG) enzymes during V(D)J recombination^5–11^. Recent studies have further highlighted the regulatory roles of 3D architectures in DNA replication^12–19^ and DNA repair^20–23^. In this context, the chromatin loop-formation and -anchoring complex Cohesin has been shown to play a critical role in DNA damage response^16,23^ and in broken end-tethering at double-stranded breaks (DSBs)^21^. Depletion of RAD21, a core subunit of the Cohesin complex, results in premature activation of dormant origins and induced replication stress in human cells^16^. On the other side, temporal chromatin structures emerging during cell cycle are also indicators of chromatin dynamics closely linked with DNA metabolism. For instance, chromatin fountains formed during DNA replication elongation reveal that replication forks remain spatially coupled from initiation to termination in mammalian cells^18^. Moreover, Cohesin-mediated stripe structures are also discovered at nascent DSBs sites, indicating the engaging of Cohesin in DNA repair^20,22^. Characterizing these highly-dynamic structures is essential for elucidating the spatial regulation mechanisms behind DNA metabolic processes.

Compared to site-specific chromatin interaction detection methods such as chromosome conformation capture (3C) or circular chromosome conformation capture (4C), the development of high-throughput chromatin conformation capture techniques (Hi-C) have provided genome-wide unprecedented insights into chromatin architecture, including A/B compartments, topologically associated domains (TADs), and chromatin loops^1,24,25^. In order to explore chromatin interactions involving specific proteins, several methods have been developed by combining chromatin interaction capture techniques with chromatin immunoprecipitation (ChIP), including chromatin interaction analysis by paired-end tag sequencing (ChIA-PET)^26,27^, Capture-C^28,29^, HiChIP^30^, and PLAC-seq^31^. These methods greatly enhance our understanding of spatial regulation of transcription, chromatin modifiers, architectural proteins and other important regulators. A recent extension of Hi-C technology, known as Replication-associated *in situ* Hi-C (Repli-HiC), specifically captures chromatin interactions near replication forks^18^. This new method has revealed chromatin structures known as chromatin fountains, which, along with stripes identified in other methods, share similar linear trajectory-like features. Despite differences in the mechanisms of their formation, these structures can be categorized as trajectory-like chromatin architectures. However, due to their highly dynamic nature, systematically resolving these chromatin trajectories and quantifying their structural variability across genomic contexts remain a significant computational challenge.

In our previous work, we developed the ‘Fun1’ to identify chromatin fountain structures across the genome, with focus on the localization of potential chromatin fountain regions^18^. Here, we introduce ‘Fun2’, an advanced computational framework for robust identification of multi-dimensional information of chromatin trajectories including both chromatin fountains and stripes, enabling a more comprehensive understanding of genome dynamics in physiological and pathological contexts.

## Results

### Overview of Fun2 framework

Trajectory-like chromatin structures, such as fountains, are widely distributed throughout the entire genome. Due to the vast size of the genome, the quantification of these structures generally involves a two-step process. The first step, known as ‘preprocessing’, focuses on the initial localization of these chromatin structures across the genome, providing crucial prior information for subsequent analyses (Fig. 1A). In the second step, termed ‘planning’, the preliminary localization is refined by adjusting the sampling box to accurately capture and align with the chromatin structure’s trajectory (Fig. 1B, C; Supplementary Fig. 1A-C).

**Figure 1.**
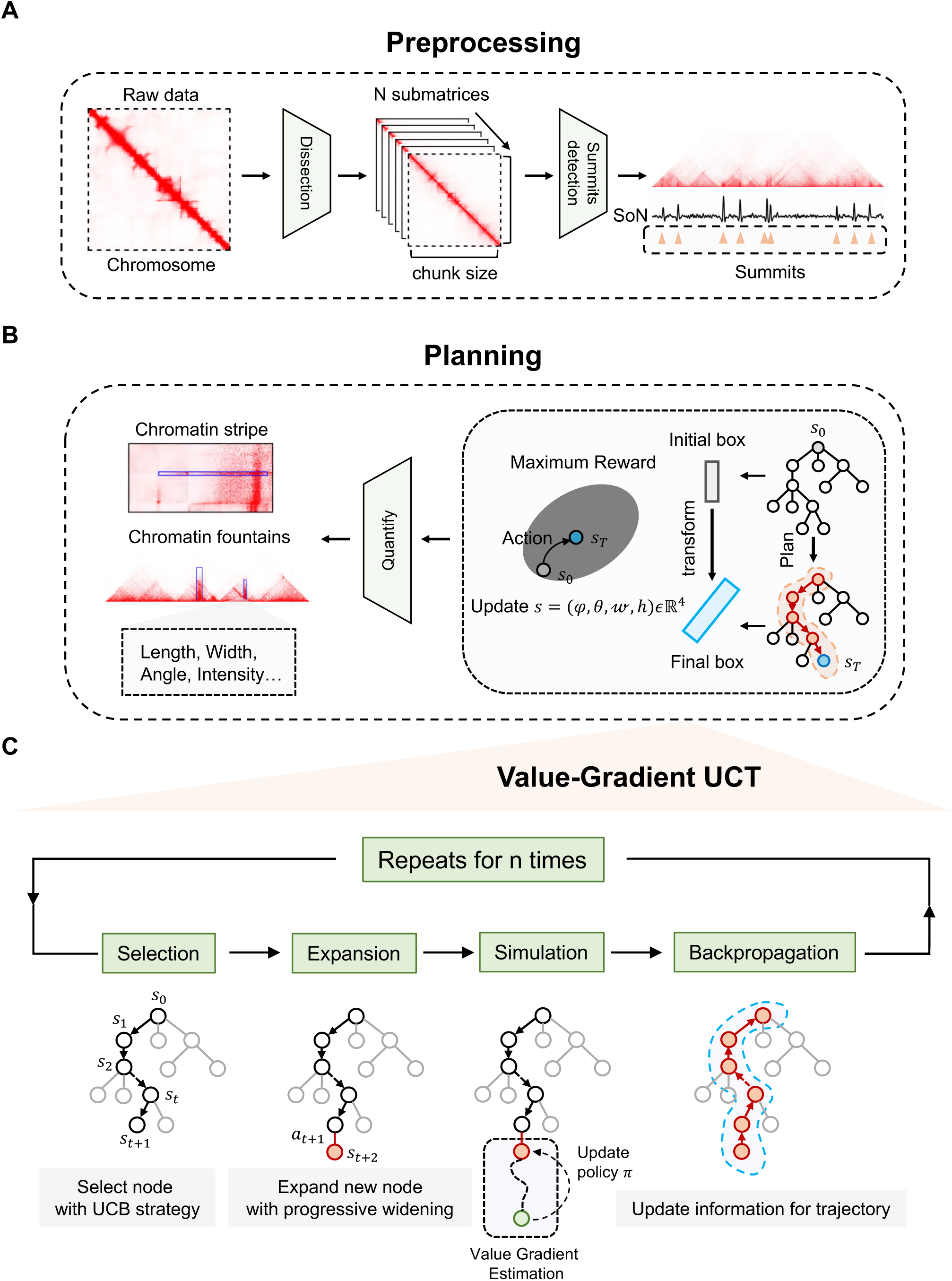
Fun2 is a reinforcement learning framework to infer trajectory of fountain-like structure. (A) Workflow for preprocessing module. The dashed line in the chromatin contact matrix indicates the regions to be processed further. Chromatin contact matrices are divided, and summits with high signal-over-noise ratios are identified as summits for subsequent analysis. (B) Workflow for planning module. The detected summits provide essential prior information for trajectory tracing, assisting in the planning of the sampling box. The VG-UCT algorithm is used to adapt the position, size, and orientation of the sampling box in this phase. The identified structures, after quality evaluation, are further quantified. (C) The details of VG-UCT algorithm. The MCTS framework consists of four main stages: selection, expansion, simulation, and backpropagation, with incorporation of the value-gradient strategy to iteratively adapt the sampling box.

In the preprocessing stage, the genome is scanned using a sampling box, with convolution operations applied to compute the signal-to-noise (SoN) metric. This metric quantifies the signal intensity within each sampling box relative to the surrounding background, effectively filtering out noise from irrelevant chromatin interactions^18^. In Fun1, a single sampling box with an initial set of parameters is used to scan the entire genome, generating a SoN profile. Regions with local maxima in the SoN signal are defined as ‘summits’, which are represented as the genomic localizations of the fountain structures. However, this approach does not fully account for the diversity of chromatin structures and potential impact of signal noise, leading to either missed detections or false-positive identifications. To overcome this limitation, Fun2 introduces several refinements. First, multiple initial sampling box parameters are used to localize and verify the summits, balancing the quantity and specificity of the detected positions (Supplementary Fig. 2). Once the sampling box parameters are roughly defined, further adjustments are made to distinguish fountain extension signals from other chromatin structures, such as chromatin loops. The summits that remain after these adjustments undergo additional thresholding, improving the specificity of these identified positions. This refined approach enables Fun2 to more robustly identify potential fountain locations during the preprocessing stage.

Building upon the preprocessing step, the ‘planning’ phase is crucial for the accurate quantification of chromatin fountains. The goal is to utilize the characteristics of local chromatin interactions at the identified summits to capture the fountain signal through the sampling box and quantify its features. In Fun1, static, fixed sampling boxes are employed, and basic numerical methods are used to analyze the characteristics of signal extension, providing a rough estimation of the fountain’s elongation length. However, this method does not account for the multi-scale variability of fountain structures, making it challenging to accurate quantification and assess their quality. In Fun2, the framework integrates Monte Carlo Tree Search (MCTS) enhanced with the Value-Gradient Upper Confidence Tree (VG-UCT) algorithm^32,33^ to trace chromatin structures (Fig. 1B, C). This integration allows Fun2 to adaptively refine the alignment of the sampling box with the chromatin structure’s trajectory by adjusting its position, size, and orientation through affine transformations (Fig. 2A; Supplementary Fig. 1C). After the planning step, the accurately positioned sampling box, along with parameters such as position, width, length, and orientation of the fountain or stripe, undergoes quality evaluation through statistical testing and scoring (Fig. 2 B-D). This ensures that only high-confidence detections are retained for further analysis (Supplementary Fig. 1D).

**Figure 2.**
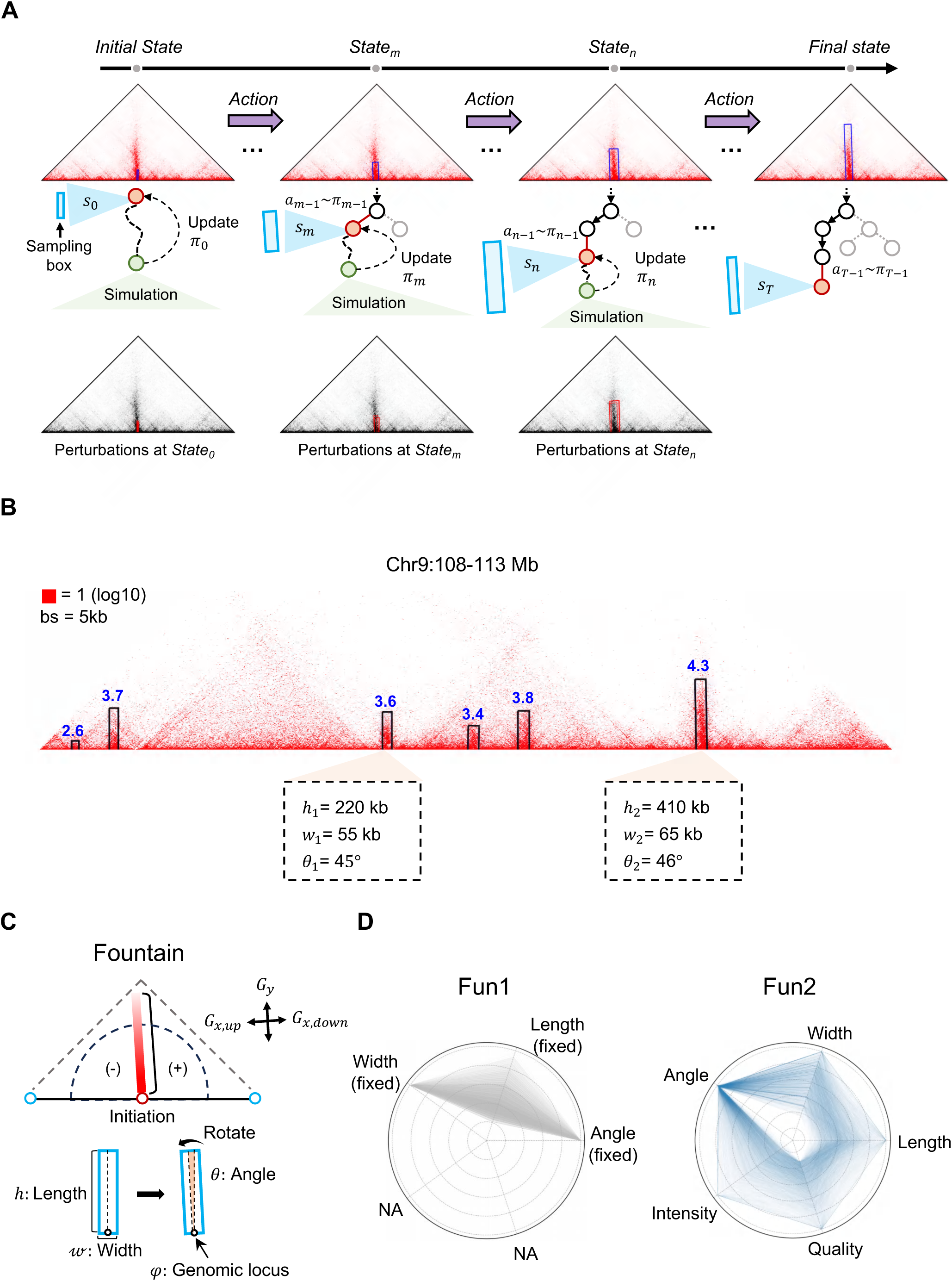
Fun2 evaluates multidimensional information of fountain. (A) An illustrative example showing the dynamic adjustment of the sampling box during MCTS. The blue box represents the sampling box, and the selected sampling box at state *s*_0_ is indicated by the red node. The green node represents the performance of the simulation. The node with a black edge, outlined by the black arrow, highlights the selected path in the search tree. The sampling box in the contact matrix shown in red color represents the dynamic adjustment process, while the grey contact matrix with red sampling box indicates the simulation process with perturbations of the sampling box. (B) An illustrative locus with multiple identified tiny-scale fountains. A locus containing multiple identified tiny-scale fountains. The black box, highlighted by blue numbers above, represents the sampling box with the corresponding fountain intensity. The yellow shaded region highlights the quantified parameters including fountain length (*h*), width (*w*) and angle (*θ*) for two typical fountains. (C) A schematic diagram defining fountain length (*h*), width (*w*) and angle (*θ*), and genomic locus (*φ*) for fountain. The black arrows represent the orientations of gradient. (D) A radar chart comparing Fun1 and Fun2. Each line represents an identified fountain. The angle, width, length, quality, and intensity are scaled from 0 to 1, with numerical values increasing as the radius extends outward.

### Benchmarking of Fun2 and Fun1

Fun2, which incorporates reinforcement learning, enables dynamic tracing of fountain trajectories starting from a sampling box with relatively loose initial parameters. This adaptive approach allows for better versatility in capturing fountains with varying characteristics across the genome. In contrast, Fun1 heavily depends on parameter tuning to identify fountains, with each set of parameters potentially yielding inconsistent results, thus limiting its flexibility for detecting both tiny and large-scale fountains (Fig. 3A, B). As a result, the quantified values for fountain length, width, and rotation angle indicate that Fun2 can capture a broad spectrum of characteristics for distinct genome-wide fountains, at both tiny and large scales. In contrast, Fun1 identifies only a subset of fountains within each resolution, constrained by the fixed parameters at each scale (Supplementary Fig. 7A-F).

**Figure 3.**
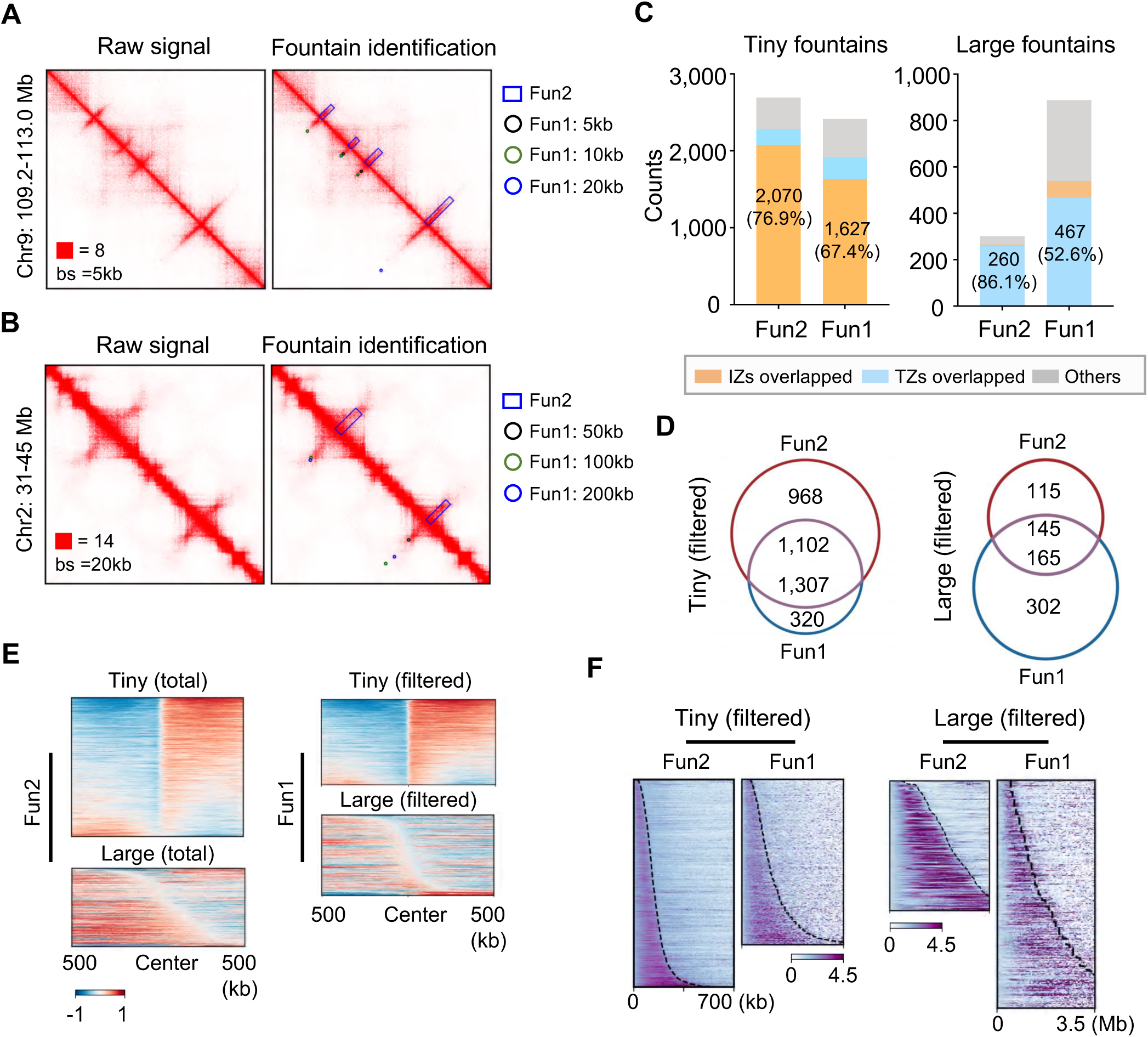
Fun2 shows more robust performance than Fun1. (A) An illustrative locus containing multiple tiny-scale fountains. Chromatin contact matrix, combined with results of identified fountains from Fun1 and Fun2, are displayed. The rectangle box with a blue edge represents the result identified by Fun2. Circles with black, green, and blue edges represent the results identified at 5-kb, 10-kb, and 20-kb data resolutions by Fun1, respectively. The red square indicates the maximum value for colormap. (B) An illustrative locus containing two typical large-scale fountains. Results are shown as described in (A). (C) The number of identified tiny-scale fountains, along with their overlap with replication initiation/termination zones. Orange box indicates the overlapped fountains with replication initiation zones (IZs). Blue box indicates the overlapped fountains with replication termination zones (TZs). Grey box indicates other fountains. (D) The Venn plot displaying the overlap between the identified tiny or large-scale fountains (after filter) by Fun1 and Fun2. (E) The heatmap showing the distribution of OK-seq signal around ±500 kb regions centered on filtered tiny or large-scale fountains identified by Fun1 and total tiny or large-scale fountains identified by Fun2. (F) The heatmap displaying the interaction signal for both filtered tiny or large-scale fountains identified by Fun1 and Fun2. Dotted lines indicate the identified end point of chromatin fountains in Fun1 and Fun2.

Replication-associated events serve as a validation metric for the identified fountains. About 220 million cis-interactions above 1-kb distance captured by Repli-HiC in K562 cells were applied for fountain identification by Fun2 and Fun1. In Fun2, 76.9% of the tiny-scale fountains exhibit clear overlap with replication initiation zones (IZs), whereas Fun1 shows a lower overlap of 67.4%. For large-scale fountains, although Fun2 identifies fewer events, it demonstrates a higher percentage (86.1%) of overlap with replication termination zones (TZs), owing to its ability to filter out summits associated with tiny-scale fountains. In contrast, only 52.6% of the large-scale fountains identified by Fun1 overlap with replication termination zones^34^ (Fig. 3C). The identified fountains—both tiny and large—are further filtered by their association with replication initiation and termination events, respectively (Fig. 3D). Pileup analysis of OK-seq signals confirms the accuracy of the filtered tiny and large-scale fountains in both Fun1 and Fun2 (Fig. 3E). Notably, limited by the lower accuracy of identified fountains by Fun1, these fountains demand further quality verification by IZs or TZs identified from OK-seq signal, while in Fun2, all the identified fountains already display high overlap ratio with replication-associated events (Fig. 3E). Moreover, when examining the heatmap of chromatin interaction profiles for the identified fountains, Fun2 reveals a distinct elongation pattern of the signal, while Fun1 shows a more ambiguous signal profile. This discrepancy suggests that the features quantified by Fun1 may be of lower quality or less accurate compared to those identified by Fun2, thus highlighting the superior performance of Fun2 in feature detection accuracy.

The quality of fountain identification is influenced by the precise placement of the sampling box, which directly impacts signal continuity and strength relative to background noise. Consequently, the fountain score (see Methods for details) serves as a robust metric for assessing the quality of fountain identification. By effectively capturing the full range of variation in fountain size—including accurate extension lengths as well as variations in width and angle (Supplementary Fig. 7A-F)—Fun2 consistently outperforms Fun1 in terms of fountain score across various resolutions. Significantly higher fountain scores reflect improved accuracy and reliability of the identification process (Supplementary Fig. 7G, H).

### Fun2 can identify chromatin stripes

Chromatin stripes, sharing similar characteristics with fountains, can be effectively identified using Fun2. During the preprocessing stage, Fun2 initializes the sampling box either horizontally or vertically, sliding it along the diagonal of contact matrices to calculate the signal-to-noise (SoN) value. Leveraging the robust search capabilities of MCTS, Fun2 iteratively refines the initial sampling box, progressively converging on the chromatin stripe (Fig. 4A; Supplementary Fig. 8A). To optimize stripe detection, a series of titration experiments for summit filtering and training iterations were conducted. A SoN threshold of 0.3 and 1,000 training iterations were found to be optimal for detecting both vertical and horizontal stripes (Supplementary Fig. 10). With this approach, Fun2 successfully traced stripe trajectories, with the width and length parameters gradually converging to their optimal values under the MCTS framework.

**Figure 4.**
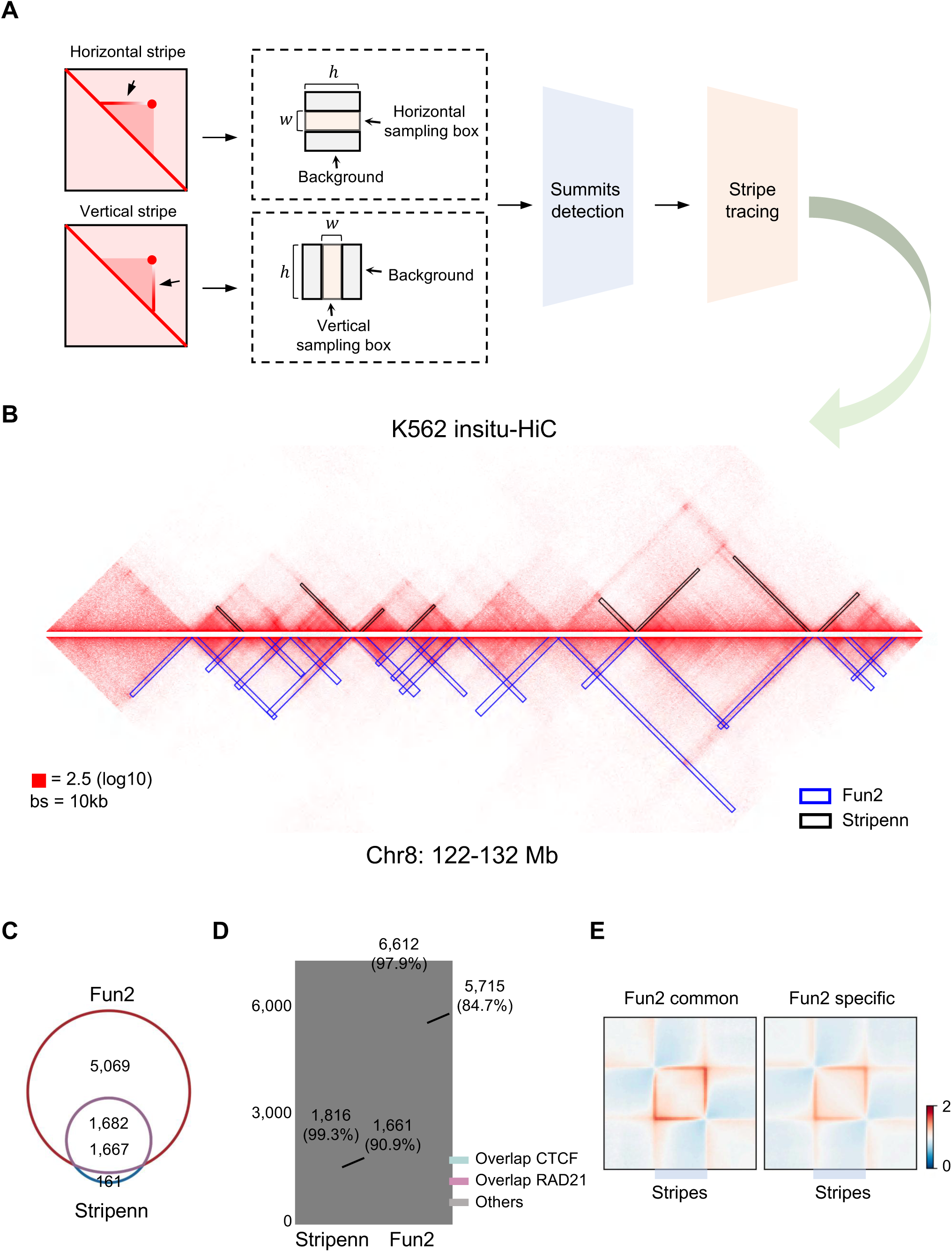
Fun2 is capable of identifying chromatin stripe. (A) Workflow of identification of chromatin stripes by Fun2. (B) An illustrative locus showing identified chromatin stripes in K562 *in situ* Hi-C data by Stripenn and Fun2, respectively. Blue box represents stripes identified by Fun2, while black box represents stripes identified by Stripenn. (C) Venn plot showing the overlap of stripes identified by Stripenn and Fun2. (D) Histogram showing the number of stripes identified by Stripenn and Fun2 and their overlap with CTCF, RAD21 enrichment sites. Pink shadow indicates the overlapped stripes with RAD21 enrichment sites. Cyan shadow indicates the overlapped stripes with CTCF enrichment sites. (E) Pileup analysis of stripes identified by both Stripenn and Fun2 (Fun2 common) or specificly identified in Fun2 (Fun2 specific) after signal rescaling.

We further compared Fun2 with ‘Stripenn’, a state-of-the-art image-based algorithm for chromatin stripe detection^35^. In K562 cells, using contact matrices with identical cis-chromatin interactions and resolution, Fun2 detected 91.2% of the chromatin stripes identified by Stripenn, while also identifying over 5,000 additional Fun2-specific stripes (Fig. 4C). Stripes identified by both Fun2 and Stripenn showed comparable positive rates, as demonstrated by the overlap ratios between stripe summits and CTCF (97.9% for Fun2 vs. 99.3% for Stripenn) and RAD21 (84.7% for Fun2 vs. 90.9% for Stripenn) enriched genomic regions (Fig. 4D). Pileup analysis further confirmed that Fun2 exhibited superior sensitivity in detecting relatively weak stripes, while maintaining robust accuracy (Fig. 4E).

## Discussion

In our analysis of chromatin fountains identified by Fun2, we have characterized five key attributes: location, intensity, length, width, and angle. Each attribute conveys distinct biological significance. Specifically, fountain location indicates the initiation or termination zones of DNA replication, aligning with the IZs identified in OK-seq and the early replication initiation zones (ERIZs) detected in NAIL-seq. Fountain intensity provides quantitative measures of specific fountains, which are crucial for analyzing fountain variations in different genotypic contexts. Fountain length reflects the distance of elongation for coupled replication forks. Notably, some fountains terminate before TZ regions in our previous analysis^18^. This phenomenon may be attributed to the limited resolution of Repli-HiC method under low signal-to-noise conditions or suggest that certain obstacles encountered during replication elongation can disrupt fork coupling without affecting the fork progression prior to replication termination^15,36^. Analysis on fountain length variations in the future may significantly advance the research on replication elongation and genome stability in mammalian cells. Fountain width exhibits enhanced flexibility and accuracy in Fun2 compared with Fun1. Specifically, Type-I fountain width accurately delineates genomic regions associated with replication initiation, whereas Type-II fountain width indicates the concentration of replication termination zones, which likely plays a crucial role in maintaining genome stability during DNA replication^18^. Additionally, while the rotation angles of fountains in Fun2 are variable, most identified fountains are nearly perpendicular to the diagonal of Hi-C 2D matrices, indicating that most coupled replication forks proceed at approximately the same rate under physiological conditions. Analyzing variations in fountain angles could provide valuable insights into the regulatory mechanisms governing the symmetry of sister replication forks. Compared to existing sequencing-based methods associated with DNA replication, the combination of Repli-HiC and Fun2 analysis substantially enriches the characterization of DNA replication features in mammalian cells simultaneously, enhancing our multidimensional understanding of DNA replication.

## Methods details

### Data preprocessing

In this study, all Repli-HiC libraries were processed using pipelines from 4DN Data Portal. Briefly, sequencing files from biological replicates were merged into a single dataset, followed by linker sequence trimming using the trimLinker tool from ChIA-PET2. The reads were aligned to the GRCh37 (hg19) genome assembly using the bwa mem algorithm. Chromatin contacts were extracted with pairtools (https://github.com/open2c/pairtools). Low-quality contacts were removed, and the remaining valid interactions were deduplicated and mapped to fragments annotated in the HaeIII-digested human genome, excluding contacts within the same fragment. Interaction distances were calculated for each contact pair, with pairs exhibiting distances smaller than 1 kb discarded. The valid contact pairs were subsequently converted into .hic and .cool formats using Juicer (https://github.com/aidenlab/juicer) and HiCExplorer (https://github.com/deeptools/HiCExplorer), respectively, and normalized using square root vanilla coverage normalization (VC_SQRT). Contact matrix at 5 kb resolution and 50 kb resolution was used to identify tiny and large fountains, respectively. For the processed K562 *in situ* Hi-C data, we used the dataset from Rao et al., 2014 (GSE63525) at a 10-kb resolution to identify chromatin stripes. The matrix was normalized by distance using observed/expected for identification.

### Initialization of sampling box and its padding background

In this study, we define the sampling box by its full state *s*, which is characterized by its genomic position *φ*, rotation angle *θ*, width *w* and height ℎ. Notably, the base of the sampling box slides along the diagonal of the contact matrix, ensuring that its position is always (*φ*, *φ*). The rotation angle *θ* defines the elongation direction of the sampling box, which plays a critical role in identifying signal gradients.

In general, a horizontally aligned sampling box (i.e., parallel to the *u*-axis of the contact matrix; Supplementary Fig. 1A) is defined with *θ* = 0°. When the box is rotated counterclockwise, the angle *θ* increases, reaching 90° for a vertically aligned sampling box (i.e. parallel to *v*-axis). For a sampling box whose elongation direction is perpendicular to the diagonal of the contact matrix, *θ* = 45°.

To describe the elongation direction more precisely, we establish a local coordinate system for the sampling box. Initially, the sampling box is set in the horizontal orientation (*θ* = 0°), where the elongation direction is defined as the *y*-axis. The direction perpendicular to the elongation is defined as the *x*-axis (Supplementary Fig. 1A).

To analyze signal gradients along the elongation direction, the sampling box is divided into *N* layers, each with a fixed height 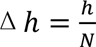. Each layer is indexed by *i*, enabling precise localization and analysis of the sampling box’s signal. The bottom layer corresponds to *i* = 1, and the top layer corresponds to *i* = *N*. Each layer of the sampling box is defined in the local (*x*, *y*)-coordinate system, where the center of the sampling box at (*φ*, *φ*) serves as the origin. The sampling region at layer *i* is defined as:

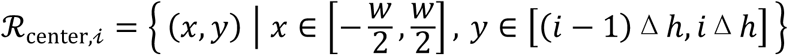

Upstream and downstream background region as:

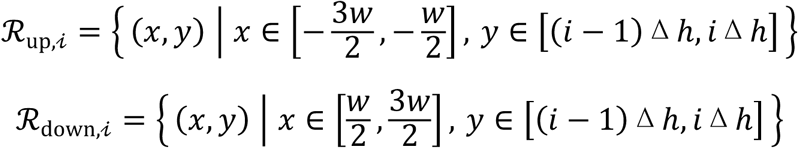

Since the contact matrix uses a global (*u*, *v*)-coordinate system, we apply a linear transformation to map the sampling box’s local (*x*, *y*)-coordinate to the global system using the following rotation matrix:

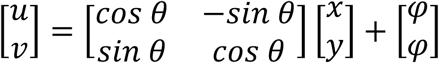

This transformation ensures that the sampling region and background regions can be consistently represented in the global (*u*, *v*)-coordinate system, allowing for precise gradient calculations.

The signal gradients along both elongation direction (*y*-axis) and the perpendicular direction (*x*-axis) are critical for adjusting the sampling box to maximize signal contrast and continuity. For each layer *i*, the upstream signal gradients are computed as follows:

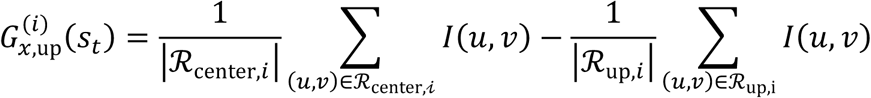

Downstream signal gradients are computed as:

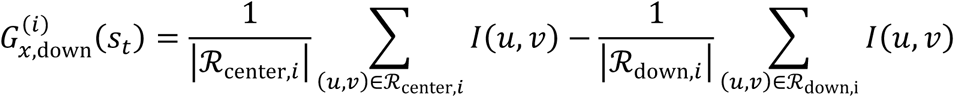

Signal gradients for the direction of elongation are computed as:

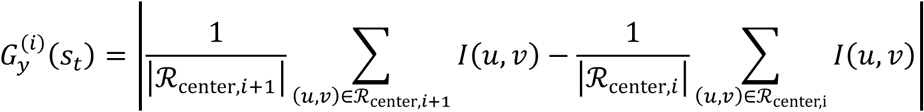

Here, *I*(*u*, *v*) denotes the signal intensity at pixel (*u*, *v*) in contact matrix, and |ℛ| is the number of pixels in the respective region.

### Quality evaluation for summits detection

In this study, the parameters width (*w*) and length (ℎ) play a critical role in influencing the capability of the Signal-to-Noise (SoN) metric, directly affecting the detection of summits. A suboptimal combination of *w* and ℎ can result in false positives or insufficient summit detection. To balance sensitivity and specificity for summit identification, we aim to optimize both parameters. The goal is to capture as many genuine summits as possible while minimizing false positives. We define the specificity *f*(*S_summits_*, *S_features_*) and sensitivity *g*(*S_summits_*, *S_features_*) as follows:

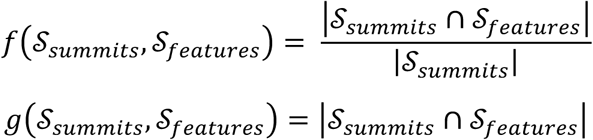

where *S_summits_* and *S_features_* represent the sets of detected summits and reference genomic features, respectively. Min-max normalization is then applied to both functions to rescale them to the [0, 1] range:

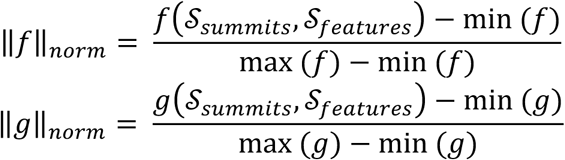

The parameters ℎ and *w* are to be optimized over their respective value ranges, ℎ ∈ ℋ and *w* ∈ *W*, by maximizing the weighted sum of the normalized specificity and sensitivity:

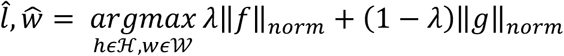

where the λ is a parameter that controls the trade-off between specificity and sensitivity. For detection of tiny-scale summits, we set ℋ = {100kb, 200kb, 300kb, 400kb, 500kb, 600kb, 700kb, 800kb, 900kb, 1000kb} and *W* = {10kb, 20kb, 30kb, 40kb, 50kb, 60kb, 70kb, 80kb, 90kb, 100kb} and used replication initiation zones for *S_features_*. For large-scale summits, we set ℋ = {1000kb, 1500kb, 2000kb, 2500kb, 3000kb, 3500kb, 4000kb} and *W* = {100kb, 200kb, 300kb, 400kb, 500kb} and used replication termination zones for validation. For stripe detection, we set ℋ = {100kb, 200kb, 300kb, 400kb, 500kb, 600kb, 700kb, 800kb, 900kb, 1000kb} and *W* = {50kb, 75kb, 100kb, 125kb, 150kb, 175kb, 200kb, 225kb, 250kb} and used CTCF-enriched sites for validation.

### Summits identification for trajectory-like chromatin structures

In our previous study, we identified fountains spanning both tiny and large scales, each predominantly associated with distinct replication events^18^. To efficiently capture these multiscale fountain structures, we employed two distinct initial sampling boxes to address the inherent variability in their scale and ensure comprehensive detection.

We conducted grid search within a defined parameter space to optimize the width and length of the sampling boxes for finding both tiny and large-scale summits (Supplementary Fig. 2A). This optimization process was guided by quality evaluation that jointly considered the number of detected summits overlapping with replication-associated events and their overlap ratios. To balance sensitivity and specificity in summit detection while minimizing noise, we introduced a hyperparameter (λ) to regulate the relative importance of these factors. For small-scale summits, the sampling box’s width was varied between 10 kb and 100 kb, and the length between 100 kb and 1,000 kb. For large-scale summits, the width ranged from 200 kb to 500 kb, and the length from 1,000 kb to 4,000 kb. Based on the balance between summit counts and overlap ratios, we found that a box with 25 kb width and 200 kb length was optimal for detecting small-scale summits, while a box with 250 kb width and 4,000 kb length was more suitable for large-scale summits (Supplementary Fig. 2B, C).

To minimize false positives arising from other chromatin structures, such as loops or stripes in fountain identification, we implemented a ‘length mutation’ strategy. Instead of using a single fixed-length sampling box, we employed three boxes of varying lengths to detect potential fountain summits. Genuine fountain structures exhibit continuous extension and consistently enriched SoN signals across all three boxes, whereas other chromatin structures like loops typically display enrichment in only one or two boxes, resulting in inconsistent SoN patterns (Supplementary Fig. 3A). This strategy effectively distinguishes genuine, high-quality loci from false-positive candidates, especially at genomic loci enriched with other types of chromatin architectures. (Supplementary Fig. 3B-E). Additionally, we applied a threshold to filter out summits with low SoN values, which are typically indicative of false-positive candidates. For tiny-scale summits, we tested SoN thresholds between 0 and 0.9 and observed that thresholds between 0.15 and 0.3 effectively balanced the number of identified summits and their overlap with replication initiation zones (Supplementary Fig. 3F). For large-scale summits, to avoid redundancy with tiny-scale summits, we excluded recurrent tiny-scale summits based on their SoN performance. We found that applying a SoN threshold between 0.2 and 0.4 can efficiently filter out approximately 80% of tiny-scale summits, while maintaining over 90% overlap ratio with replication termination events (Supplementary Fig. 3G, H).

Summit detection provides critical prior information for the initialization of the sampling box at the candidate summits. The primary objective following summit detection is to adjust the initialized sampling box so that it aligns with the fountain’s signal. In contrast to the static sampling box in Fun1, Fun2 transforms the sampling box into a dynamic, adjustable entity. As a result, while the initial locus of the sampling box is determined by the position of the candidate summit, it is not fixed. Instead, the box can autonomously trace the fountain’s trajectory, iteratively refining its orientation, width, length, and position through affine transformations, progressively aligning with the fountain structure under MCTS guidance (Fig. 2A). This flexibility allows the box to adapt to the varying shapes of fountains across different genomic loci (Fig. 2B). The parameters of the sampling box are interdependent, meaning that inaccuracies in one parameter can impact the others, leading to misalignment and preventing accurate tracing of the fountain structure. This highlights the importance of simultaneously optimizing all parameters to capture the characteristics of the fountains. In this regard, Fun1, being static, can only identify the length of the fountain within a fixed range, thus not accounting for these interdependencies. In contrast, Fun2 can quantify multiple parameters, including width, length, quality, intensity, and rotation angle, exanimating the spatial organization of DNA replication more comprehensively. This ability not only holds potential biological significance but is also crucial for technical precision, ensuring the accuracy of fountain identification (Fig. 2C, D).

To assess the stability and accuracy of Fun2 under varying numbers of training iterations, we conducted experiments with MCTS training iterations ranging from 25 to 2,000 times, with three repetitions at each experiment (Supplementary Fig. 4A). For tiny-scale fountains, we found that when the number of training iterations exceeded 250 times, the variance in the fountains’ quality scores were stabilized, and the variance of top 50% of high-quality fountains remained consistent across all training sessions, as exemplified of the fountains in chromosome 20 (Supplementary Fig. 4D, E). Averaging over the three repetitions effectively enhanced the stability of fountain quality (Supplementary Fig. 4B, C, F, G). Similar results were observed for large-scale fountains (Supplementary Fig. 5), with the quality score variance stabilizing at 250 training iterations (Supplementary Fig. 5B). As a result, at least 250 training iterations are needed for stable and accurate fountain identification with Fun2.

To evaluate the performance of Fun2 across varying numbers of chromatin interactions, we conducted a downsampling analysis ranging from 20% (approximately 40 million cis-chromatin interactions) to 100% of the total cis-chromatin interactions (approximately 210 million cis-chromatin interactions) of Repli-HiC library. We observed that at lower level of chromatin interactions, the SoN values fluctuated significantly, resulting in increased ratio of false-positive summits. As the number of interactions increased, these meaningless fluctuations diminished, while maintaining accurate identifications of genuine summits (Supplementary Fig. 6A). Consequently, the total number of identified summits gradually decreased, with the ratio of genuine summits increased as the chromatin interactions increased from 20% approached to 100% (Supplementary Fig. 6B, C). Furthermore, the quality of the identified fountains remained stable, demonstrating that Fun2 performs robustly and reliably at genuine fountain loci (Supplementary Fig. 6D).

As for the summit identification for chromatin stripes, we employed an initial sampling box with a length range of 100-1,000 kb and a width range of 25-250 kb, evaluating quality by balancing overlap counts and ratios using CTCF-enriched regions to detect both vertical and horizontal stripes. For all λ values, a sampling box with a length of 900-1,000 kb proved to be optimal, while the optimal width varied between 25 kb and 250 kb depending on λ (Supplementary Fig. 9B, C). Since wider sampling boxes may encompass more CTCF-enriched sites, potentially inflating performance metrics, we further validated the SoN ratio for boxes with a fixed length of 1,000 kb and varying widths (25 kb, 50 kb, 225 kb, and 250 kb). The results showed that boxes with widths of 25 kb and 50 kb provided more precise summit detection compared to wider boxes (Supplementary Fig. 10A, B).

### MCTS planning for identified summits

In this study, Fun2 utilizes MCTS to optimize the placement and transformation of a sampling box on an image to maximize its alignment with both fountain and stripe signal. MCTS is a simulation-based planning framework widely used in sequential decision-making problems formulated as a continuous Markov Decision Process (MDP)-a popular paradigm in reinforcement learning. In an MDP, the environment is defined by ℳ = (*S*, 𝒜, 𝑃, 𝑟, 𝛾), where:

State space *S*: The state at time step 𝑡, denoted by *s_t_*𝜖*S* ⊂ ℝ^4^, represents the sets of parameters of the sampling box, which includes its genomic locus *φ*, rotation angle *θ*, width *w* and height ℎ, which have been described above. Formally, *s_t_* = (*φ_t_*, *θ_t_*, *w_t_*, ℎ*_t_*).

Action space 𝒜: The action space 𝒜 ⊂ ℝ^4^ represents the affine transformations that can be applied to the sampling box, Specifically, at time step 𝑡, the action 𝑎*_t_* is a set of actions in continuous space and represented as: 𝑎*_t_* = (Δ*φ_t_*, Δ*θ_t_*, Δ*w_t_*, Δℎ*_t_*). The affine transformation generally includes translocation, rotation, expansion, extension along the diagonal of the contact matrix.

Transition probability 𝑃: For the sampling box at time step 𝑡 with state *s_t_*, the transition function 𝑃(*s_t_*_+1_|*s_t_*, 𝑎*_t_*) indicates the transition probability of reaching state *s_t_*_9:_ by taking affine transformation 𝑎*_t_*. In addition, 𝑃(𝑟*_t_*_+1_|*s_t_*, 𝑎*_t_*) gives the probability of receiving reward 𝑟*_t_*_+1_ after taking action 𝑎*_t_*. In our study, both the state and reward transition are deterministic.

Reward Function 𝑟: The 𝑟(*s_t_*, 𝑎*_t_*) 𝜖ℝ indicates the immediate reward at time 𝑡 given state *s_t_* and action 𝑎*_t_*. It evaluates how well the sampling box aligns with the trajectory of a fountain or stripe, generally comprising two components: (1) Maximizing contrast between the sampling box and its surrounding background. (2) Capturing as much signal as possible along the trajectory’s elongation direction. By maximizing the image gradient field, Fun2 transforms the trajectory-tracing problem of chromatin fountains or stripes into an optimization problem. Concretely, the reward is defined as:

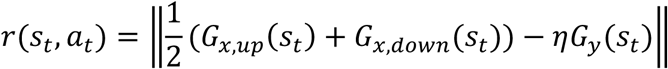

where 𝐺*_x,up_*(*s_t_*) and 𝐺*_x,down_*(*s_t_*) represent gradients in the upstream and downstream backgrounds, respectively. 𝐺*_y_*(*s_t_*) is the gradient along the elongation direction. 𝜂 indicates the hyperparameter that controls the penalty for over-elongation of the sampling box. Specifically, the 𝐺*_x,up_*(*s*), 𝐺*_x,down_*(*s*) and 𝐺*_y_*(*s_t_*) could be defined as: 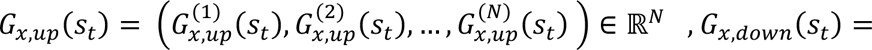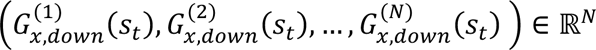 and 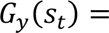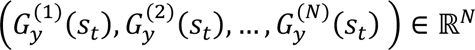.

Discount Factor 𝛾 : In the MDP, the discount factor 𝛾 controls how future rewards are weighted relative to immediate rewards. In this study, we currently use 𝛾 = 1, emphasizing immediate rewards rather than long-term returns.

Policy 𝜋: A policy 𝜋 is a mapping from states to actions, defined as 𝜋: *S* → 𝒜: where 𝜋*_t_* indicates the probability of selecting action 𝑎*_t_* at state *s_t_*, and could be represented as 𝜋(𝑎*_t_*|*s_t_*).

By incorporating MCTS, Fun2 maps the decision process onto a search tree, where nodes represent the state of the sampling box at each time step, and edges represent the actions that transition one state to another. This setup allows Fun2 to iteratively refine the sampling box to solve an optimization problem, ultimately converging on a best state that models either a fountain or a stripe configuration.

MCTS typically comprises four main steps: selection, expansion, simulation, and backpropagation, each designed to incrementally build and explore a search tree while balancing exploration and exploitation.

Selection: The selection phase traverses the existing search tree from the root node (representing *s*_0_) to find the most promising child node. This process uses the Upper Confidence Bound (UCB)^37^ strategy to balance exploration (visiting less-explored nodes) and exploitation (focusing on nodes with high rewards). Formally, the chosen action 𝑎*_t_* for a state *s_t_* is:

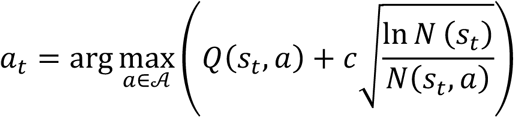

where 𝑄(*s_t_*, 𝑎) is the average reward for action 𝑎 in state *s_t_*. *N*(*s_t_*) is the number of times that state *s_t_* has been visited. *N*(*s_t_*, 𝑎) is the number of times action 𝑎 has been taken from state *s_t_*. The constant 𝑐 adjusts the trade-off between exploration and exploitation.

Expansion: If the selected node is not fully expanded (i.e., not all possible actions have been explored), a new child node is created by sampling a new action 𝑎*_t_* from policy 𝜋*_t_* and applying it to the current state *s_t_*. To handle continuous action space, we adopt ‘Progressive Widening’^38,39^, where a new child node is added only when:

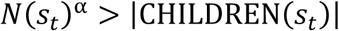

where α ∈ (0,1) controls the growth rate, and *N*(*s_t_*) represents the visit count for state *s_t_* and |CHILDREN(*s_t_*)| denotes the number of child nodes of state *s_t_*. This strategy dynamically controls the growth of the search tree in large or continuous action spaces. It prevents computational overload by incrementally adding new child nodes only when the visit count of the parent state increases sufficiently. This ensures efficient exploration of the action space while avoiding unnecessary expansion of low-reward nodes.

Simulation: During simulation, a rollout estimates the performance of the newly expanded node. These simulated data can be utilized to effectively refine the strategy 𝜋*_t_*, guiding future action generation. Here, we adopt Value-Gradient UCT (VG-UCT) algorithm^32^, which leverages first-order gradient information of reward with respect to the action to refine 𝑎*_t_*. This approach significantly improves precision and efficiency over standard MCTS methods relying purely on random rollouts. We additionally incorporate Evolution Strategies (ES) to approximate the gradient of the reward function in continuous, high-dimension al state-action spaces^40^. Specifically, the steps are as follows:

First, sample noise vectors:

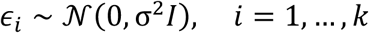

where 𝑘 is the total number of samples, 𝜎 is the standard deviation, and *I* is the identity matrix. Then, for each noise sample 𝜖_1_, compute the perturbed action 𝑎*_t_* + 𝜖_1_ and evaluate its reward 𝑟(*s_t_*, 𝑎*_t_* + 𝜖*_i_*). Using finite differences, estimate the gradient of the reward function with respect to the action at step time 𝑡:

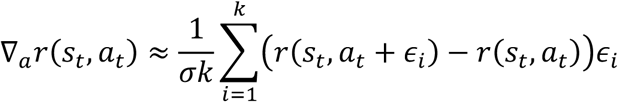

Then, adjust the action based on the estimated gradient incorporated with momentum:

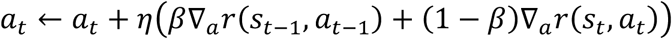

where 𝛽 ∈ [0,1] is the momentum coefficient, which determines the contribution of the previous gradient. ∇*_a_*𝑟(*s_t_*, 𝑎*_t_*) represents the gradient of the reward function with respect to the action at the current state *s_t_*. ∇*_a_*𝑟(*s_t_*_-1_, 𝑎*_t_*_-1_) is the gradient of the reward function respect to the action at the previous state *s_t_*_-1_. Further, we use dynamic action clipping to refine the step for updating actions. For example, for actions at step time 𝑡 : 𝑎*_t_* = (Δ*φ_t_*, Δ*θ_t_*, Δ*w_t_*, Δℎ*_t_*), the clipped actions can be defined as follows:

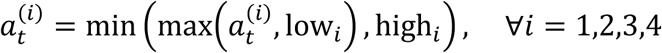

where 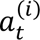 represents the *i*-th component of the action vector 𝑎*_t_*, low*_i_* and high*_i_* represent the lower and upper bounds for the *i*-th dimension of the action space. This formula ensures each component of the action vector is constrained to stay within the feasible range defined by its respective bounds.

### Quantification and quality evaluation for identified structure

For the identified fountain or stripe, Fun2 characterize multiple dimensional features, including genomic positions (*φ*), extension length (ℎ), width (*w*), rotation angle (*θ*) and signal intensity. The position is defined by the sampling box’s center in the genome, while the width and height are determined from *w* and ℎ of sampling box, The rotation angle *θ* follows the definitions above and is updated according to the sampling box’s orientation. Signal intensity is represented by the sum of chromatin interactions within the sampling box and further processed by log transformation. Let the final state of the sampling box be denoted as 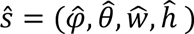, where 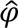 is the estimated genomic position, 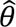 represents the rotation angle, 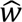 is the estimated width, 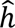 the height. The height 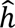 and width 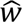 are defined under *θ* = 0°. We apply a transformation to obtain the rotated height and width:

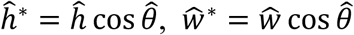

Thus, the transformed state of the sampling box, accounting for rotation, is given by:

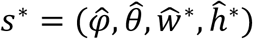

In addition, Fun2 perform quality evaluation for positioned sampling box. This involves calculating the quality score and providing statistical significance by comparing signal within the sampling box to that in local background. Specifically, the quality score, inspired by the work of Yoon et al., (2022)^35^, and is determined by two key factors: (1) the strength of the sampling box’s internal signal, and (2) how the signal behaves relative to the background in upstream or downstream regions. If the final state of sampling box is *s*^∗^, the quality score cis defined as:

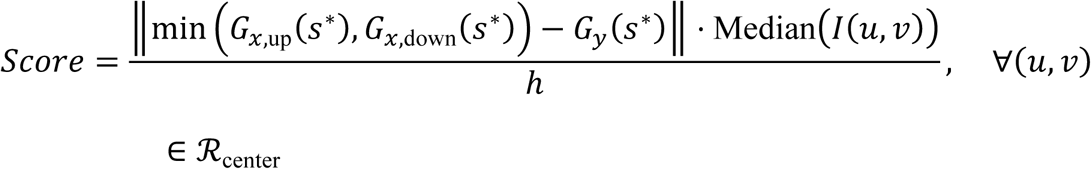

where 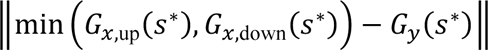 reflects the how well the signal in sampling box distinguishes from the background, and MedianQ*I*(*u*, *v*)R represents the overall signal strength inside the sampling box. The score is further normalized to the extension length of fountain or stripe, considering the inherent variability in extension length. Higher scores indicate clearer extensions and stronger signal-to-noise ratios, while lower scores suggest ambiguous or weak features.

Fun2 also performed statistical analysis to assess the final position of the sampling box. Specifically, one-sided Wilcoxon test is applied to compare the signal intensity for pixels of the sampling box (for every pixel(*u*, *v*) ∈ ℛ_center_) to the upstream (for every pixel (*u*, *v*) ∈ ℛ_up_) or downstream (for every pixel (*u*, *v*) ∈ ℛ_dowm_) background. This approach uses local background comparison and avoids assuming a specific distribution for the data. The Benjamini-Hochberg correction is used to adjust for any false discovery rate (FDR) issues. The final FDR values, combined with quality score are then used to assess the quality of identified fountains or stripes.

## Author contributions

J.H. supervised the project; Z.Z. designed the Fun2 algorithm and analyzed the data; X.L. generated the Repli-HiC library. Z.Z., X.L. and J.H. wrote the paper.

---

**Supplementary Figure 1.**
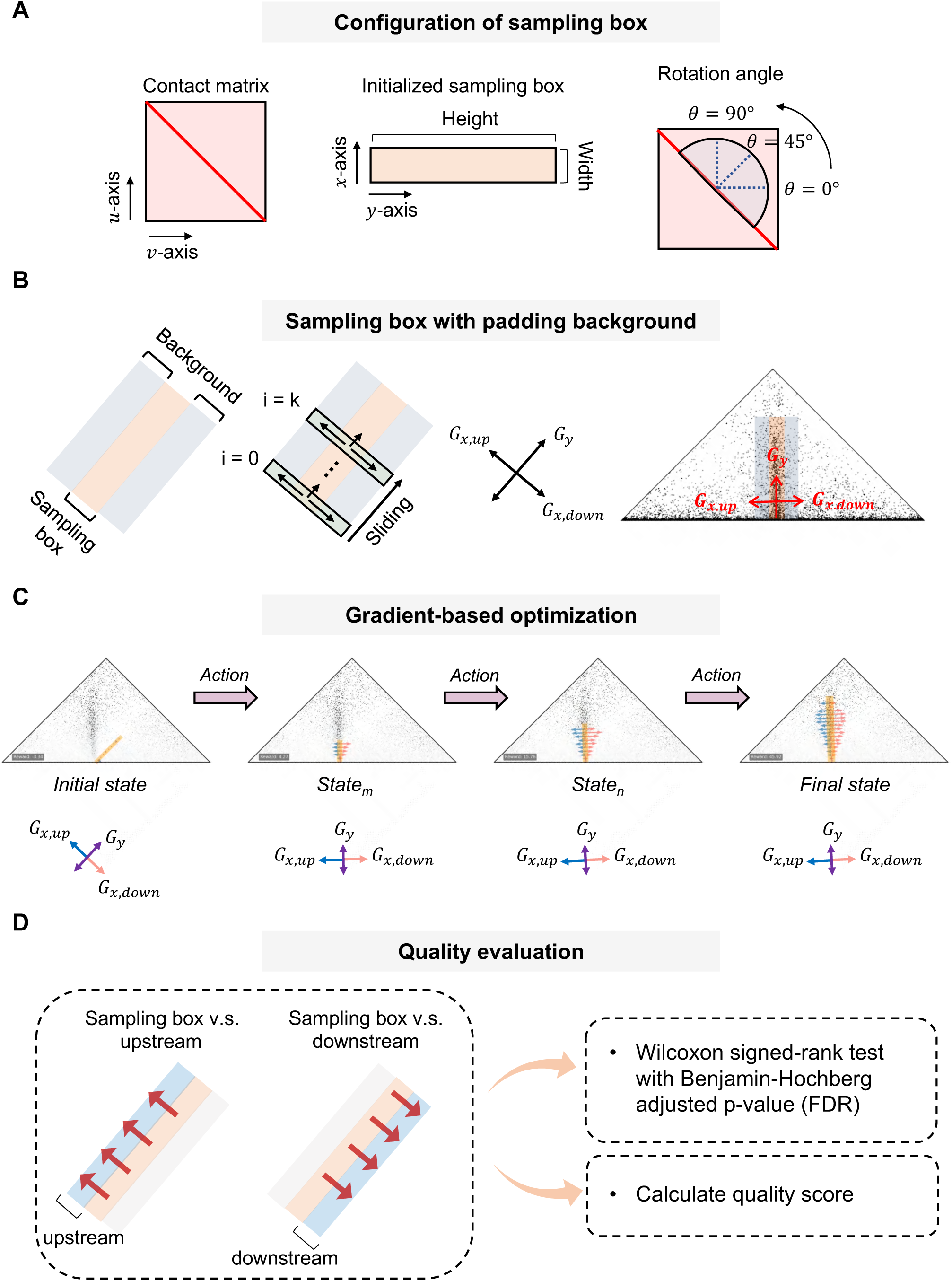
(A) Schematic representation of the sampling box and its coordinate system. (B) Composition of the sampling box and its background. The yellow region represents the sampling box, which is the area used to collect signals. The grey regions at upstream or downstream indicate the background areas, which are used to assess the signal enrichment in the central sampling area. The direction of the arrows marks the coordinate system of the sampling box, which the x and y directions indicated. On the right of the panel, the sampling box and its background regions are annotated on a fountain site, with red arrows indicating the direction of the gradient along the fountain’s extension (*𝐺_y_*) and the gradient of the central area relative to the background (*𝐺_x,up_* and *𝐺_x,down_*). (C) Iterative sampling box adjustment for fountain trajectory optimization. The sampling box dynamically updates its position and orientation to converge on the fountain trajectory by solving an optimization problem based on the local gradient field of the image. At each iteration, the box evaluates gradients in its local *x*- and *y*-coordinate system to determine the optimal position. The changes in the gradient directions (𝐺*_y_*, *𝐺_x,up_* and *𝐺_x,down_*) during the optimization process are highlighted in purple, blue and light red, respectively, and the local coordinate axes of the sampling box are continuously updated throughout the procedure. (D) Quality evaluation of the sampling box. The blue region and red arrows indicate the comparison between the sampling box and the upstream and downstream areas to assess the signal enrichment quality in the central region. Specifically, the center and background signals are evaluated using one-sided Wilcoxon test, and the quality score is calculated. Both metrics are used to assess the quality of the fountains.

**Supplementary Figure 2.**
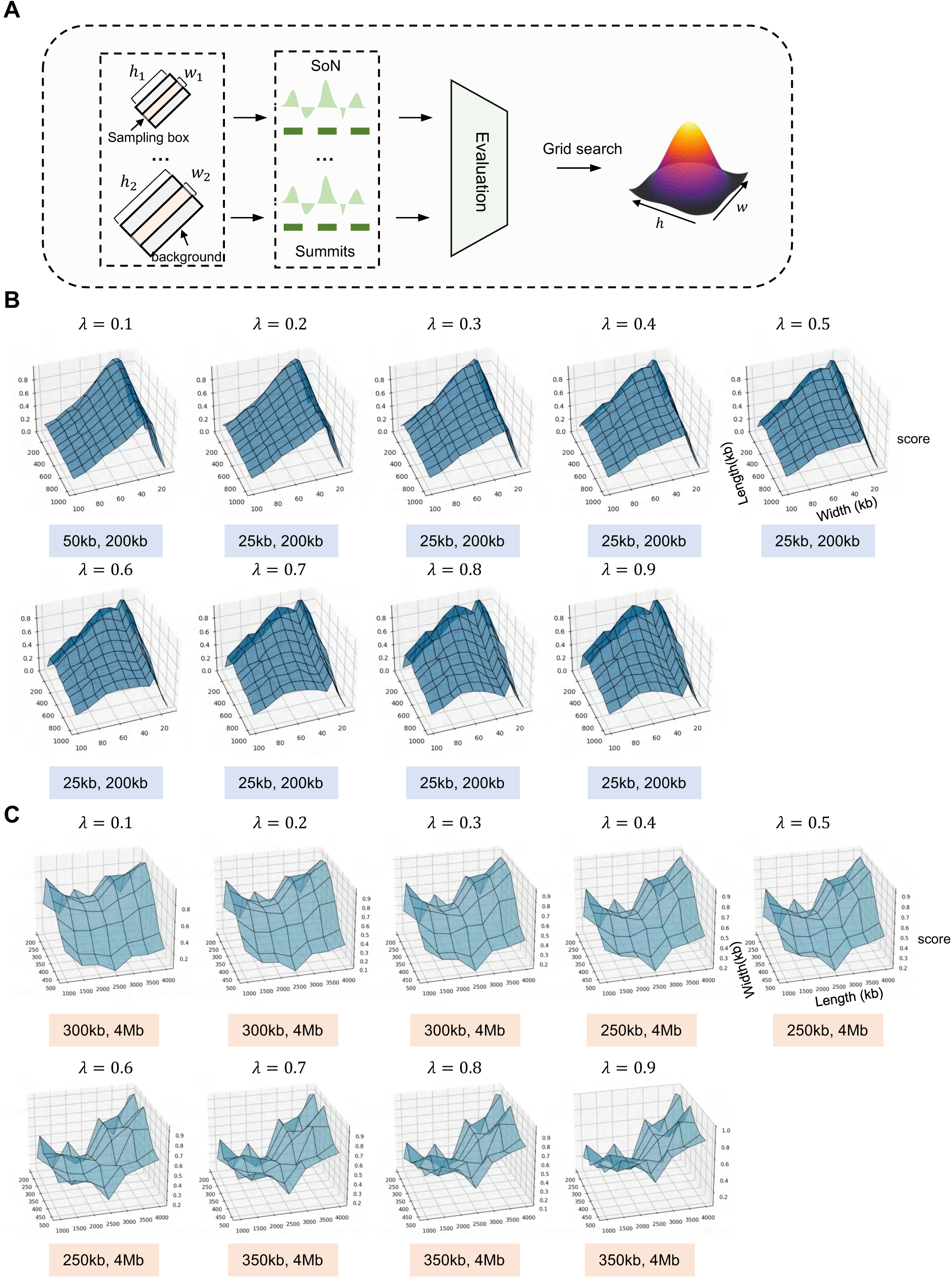
(A) Schematic of sampling box adjustment for summits detection. Using grid search, the height and width of the sampling box are adjusted within a specific range. The SoN values are calculated for each parameter combination, different parameter combinations are evaluated based on the assessment criteria. (B) Grid search results for tiny-scale summits. Result of grid search for tiny-scale summits, with hyperparameter λ ranging from 0.1 to 0.9 is shown. For each λ, the length and width of the sampling box are adjusted within a defined range, as represented by the x and y axes in 3D plot. The z-axis of the 3D plot represents the score, which is used to evaluate the quality of the identified summits for each combination. The light blue box indicates the parameters with the highest score. (C) Grid search result for large-scale summits. The process is the same as in (B), but applied to the evaluation of large-scale summits. The yellow box indicates the parameters with the highest score.

**Supplementary Figure 3.**
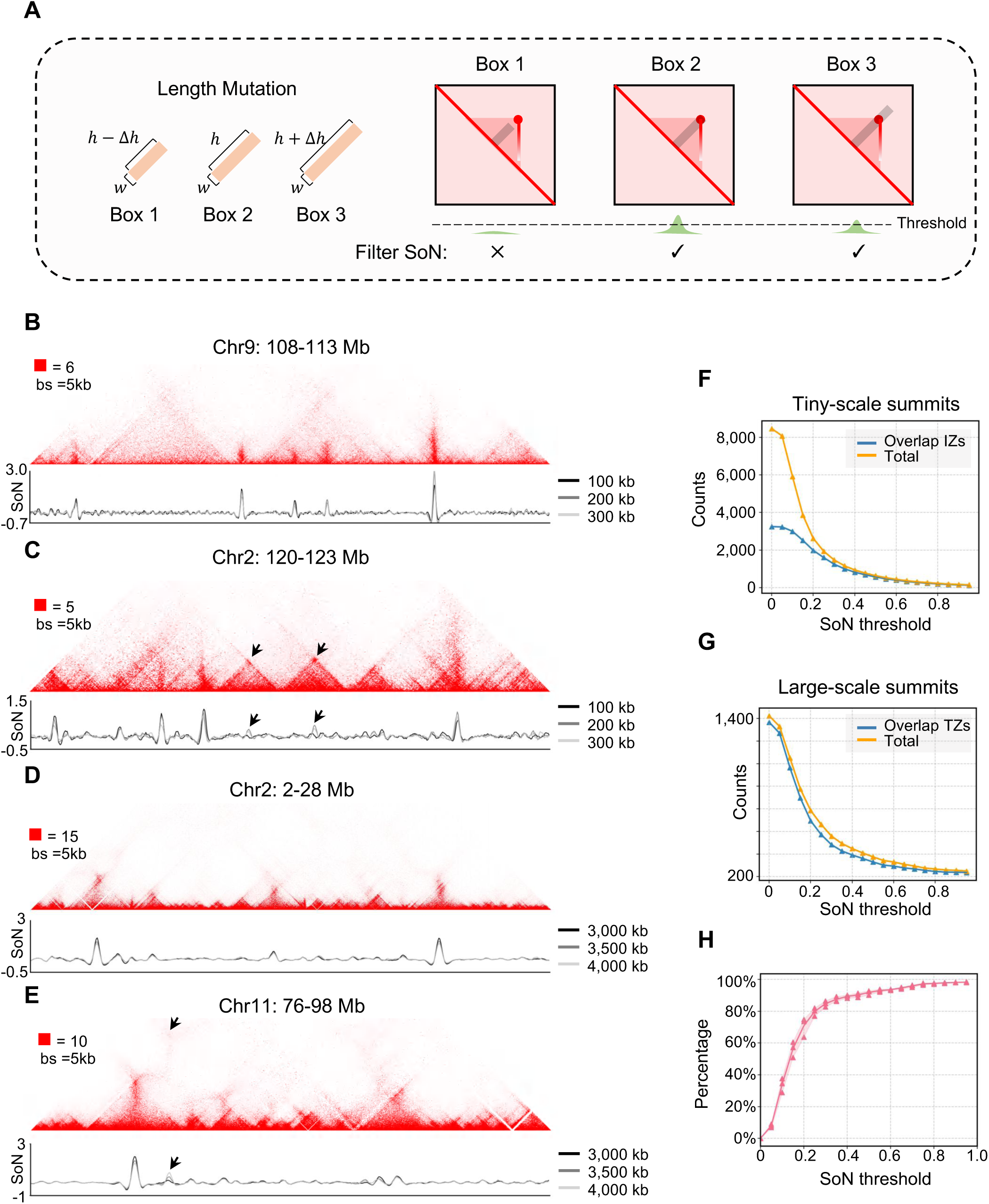
(A) Schematic of ‘length mutation’ strategy. Sampling box of varying length, with a fixed width, are used to calculate SoN signal. Different box lengths (long, medium, or short) effectively distinguish genuine chromatin fountains from other chromatin architectures. (B) An illustrative locus on Chr9 showing clear tiny-scale chromatin fountains, with the corresponding SoN signal plotted below. The black, grey, and silver lines represent SoN curves identified with boxes of indicated lengths. (C) An illustrative locus on Chr 2 showing both tiny-scale chromatin fountains and other chromatin architectures, such as loops and stripes (indicated by black arrows). The SoN values are plotted below. (D) An illustrative locus on Chr2 with large-scale chromatin fountains, with SoN signal plotted below. (E) An illustrative locus on Chr11 showing both clear large-scale chromatin fountains and other chromatin architectures (indicated by black arrows). (F) Number of tiny-scale summits at different SoN thresholds. The blue line indicates the number of summits that overlap replication initiation zones at varying SoN thresholds (Overlap IZs), while the orange line shows the total number of summits at each SoN threshold (Total). (G) The same analysis as in (F), applied to large-scale summits. (H) Percentage of summits retained at various SoN thresholds for those identified under both large-scale and tiny-scale conditions.

**Supplementary Figure 4.**
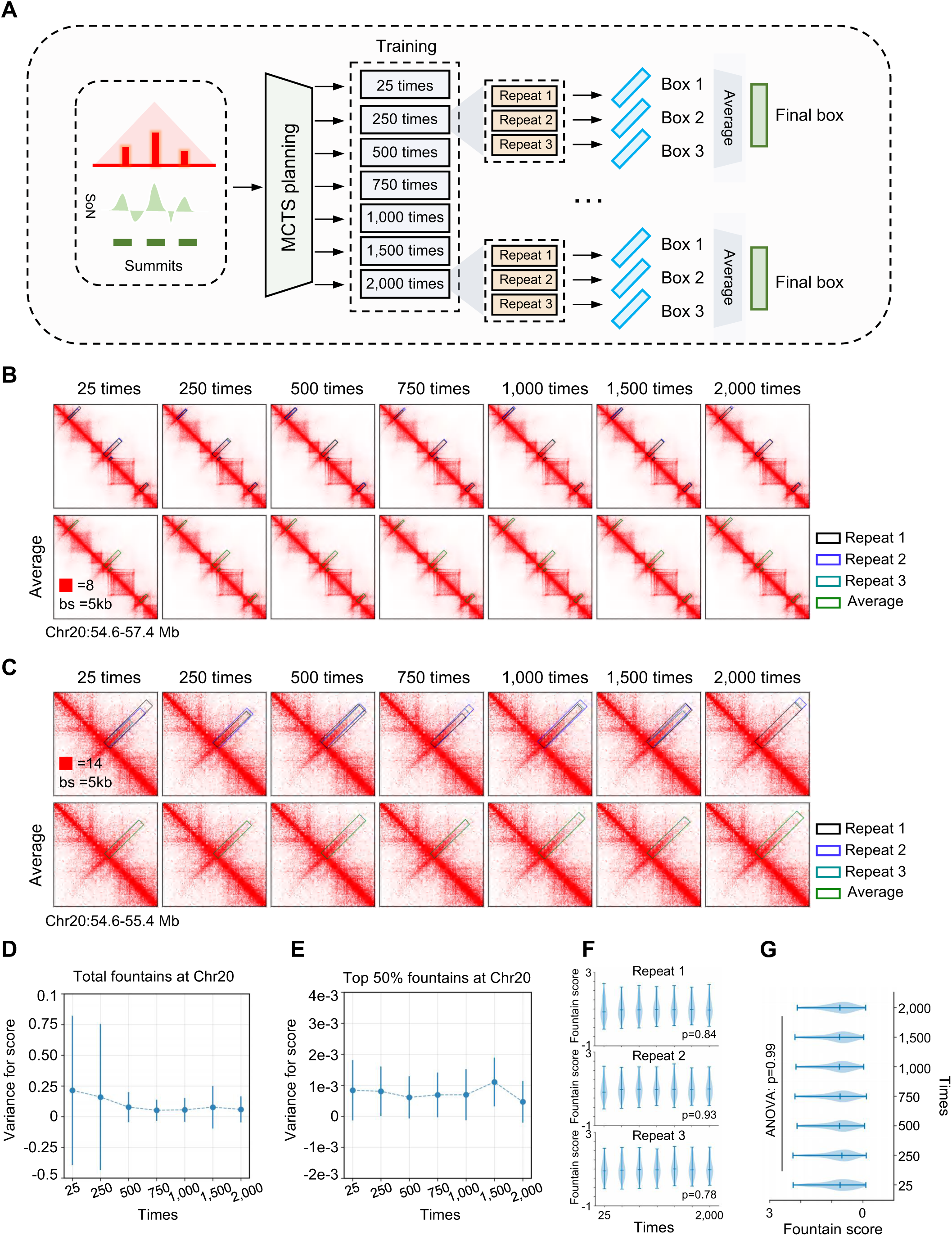
(A) Workflow of titration for MCTS training. Identified summits were submitted to MCTS training for 25 to 2,000 iterations. Three repeats were performed for each training, and the average features of the three repeats are displayed and regarded as final box. (B) An illustrative locus on Chr20 showing multiple tiny-scale fountains. The identified sampling box for repeats 1, 2, and 3, as well as the average result, are displayed with box with black, blue cyan, and green edges, respectively. (C) A zoom-in view of a typical tiny fountain displayed in (B). (D) The variance in quality score of tiny fountains on Chr20 across different training times, based on the average result. (E) The variance in quality score for the top 50% of tiny fountains (average result) at different training times. (F) The quality score for tiny fountains on Chr20 identified at repeats 1, 2, and 3 at different training times, with statistical testing performed by One-way ANOVA. (G) The quality score for fountains identified from the average results on Chr20, with statistical testing performed by One-way ANOVA.

**Supplementary Figure 5.**
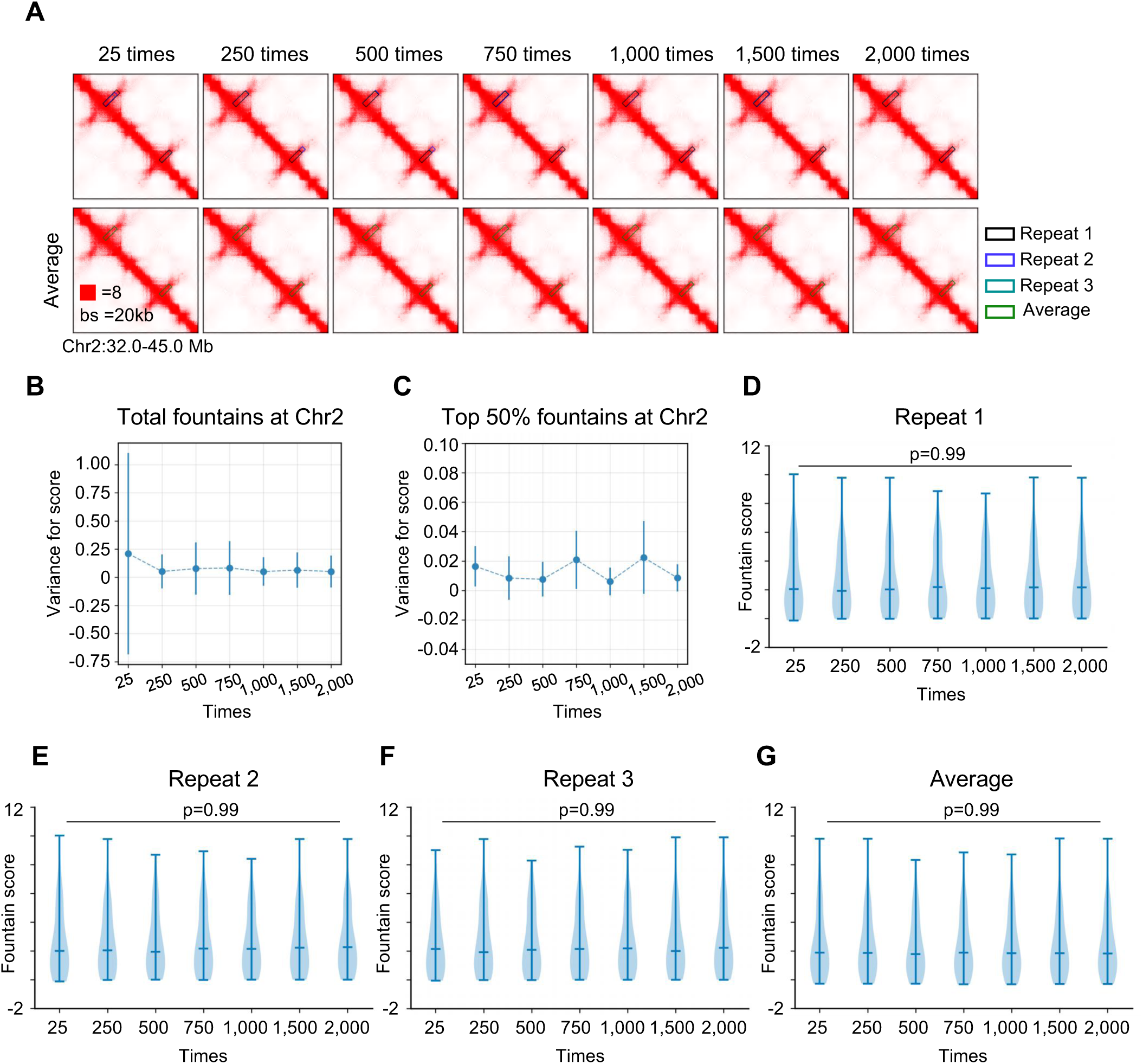
(A) An illustrative locus on Chr2 showing multiple large-scale fountains. The identified sampling box for repeats 1, 2, and 3, as well as the average result, are displayed with box with black, blue cyan, and green edges, respectively. (B) The variance in quality score of large-scale fountains on Chr2 across different training times, based on the average result. (C) The variance in quality score for the top 50% of large-scale fountains (average result) on Chr2 across different training times. (D) The quality score for large-scale fountains identified at repeats 1 at different training times, with statistical testing performed by One-way ANOVA. (E) Same as (D) but for repeat 2. (F) Same as (D) but for repeat 3. (G) The quality score for large-scale fountains identified at repeats 1, 2, and 3 at different training times, with statistical testing performed by One-way ANOVA.

**Supplementary Figure 6.**
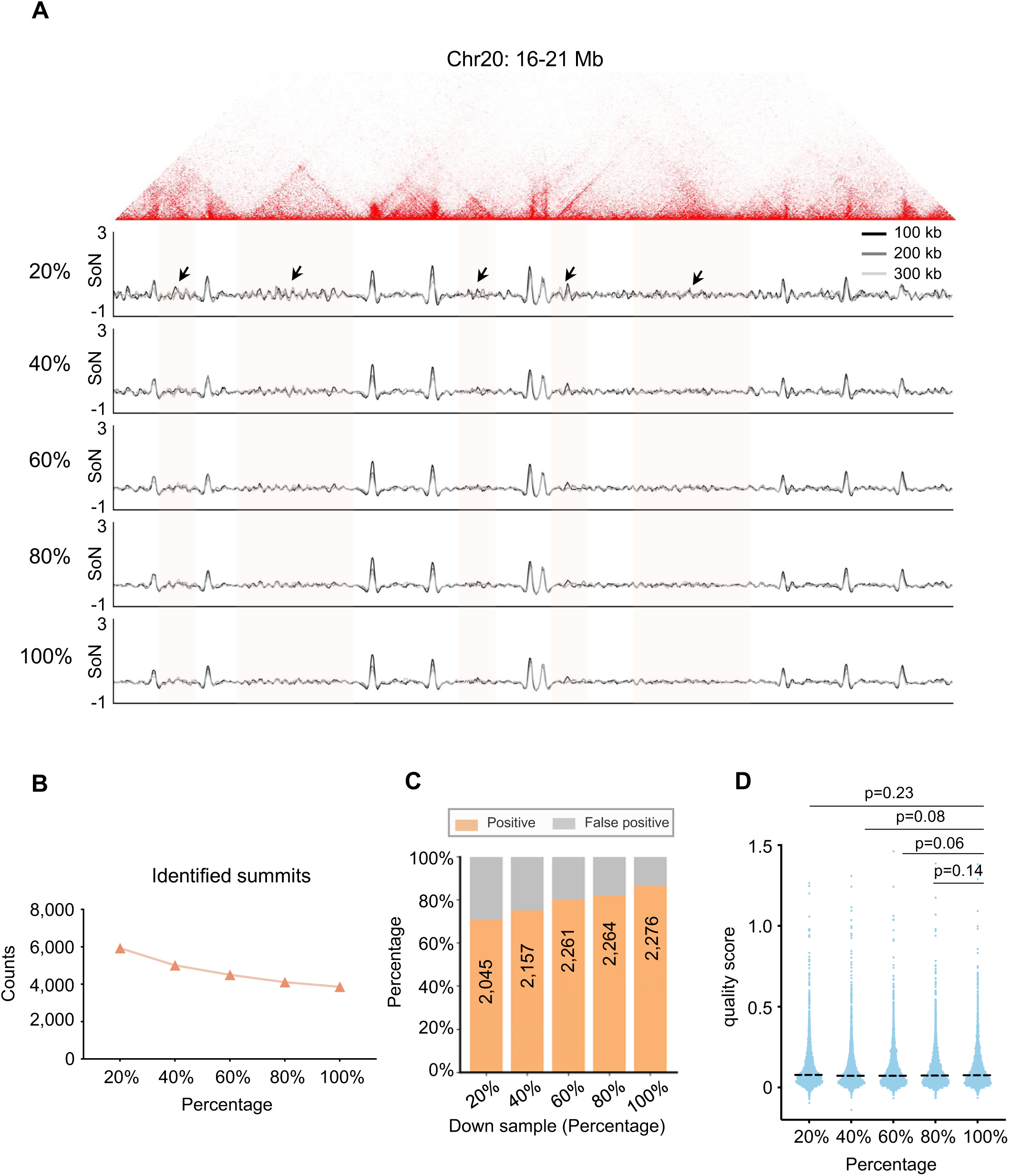
(A) An illustrative locus on Chr20 displaying Repli-HiC and its signal-over-noise (SoN) at varying numbers of cis-chromatin interactions, downsampled from 20% to 100%. Of note, 100% represents 220 million cis-interactions above 1kb distance. SoN, calculated by a sampling box with fixed width and varying lengths of 100 kb, 200 kb, and 300 kb, is represented by black, grey, and silver lines, respectively. The yellow shaded regions with black arrows indicate false-positive summits that could randomly emerge due to limited data. (B) The number of summits identified at different downsampling levels. (C) The rate of summits overlapped with IZs or TZs (positive) identified at different downsampling levels. Number of positive summits are labeled on the chart. (D) The distribution of quality scores for fountains identified at different downsampling levels. Statistical significance was assessed using one-sided Wilcoxon test.

**Supplementary Figure 7.**
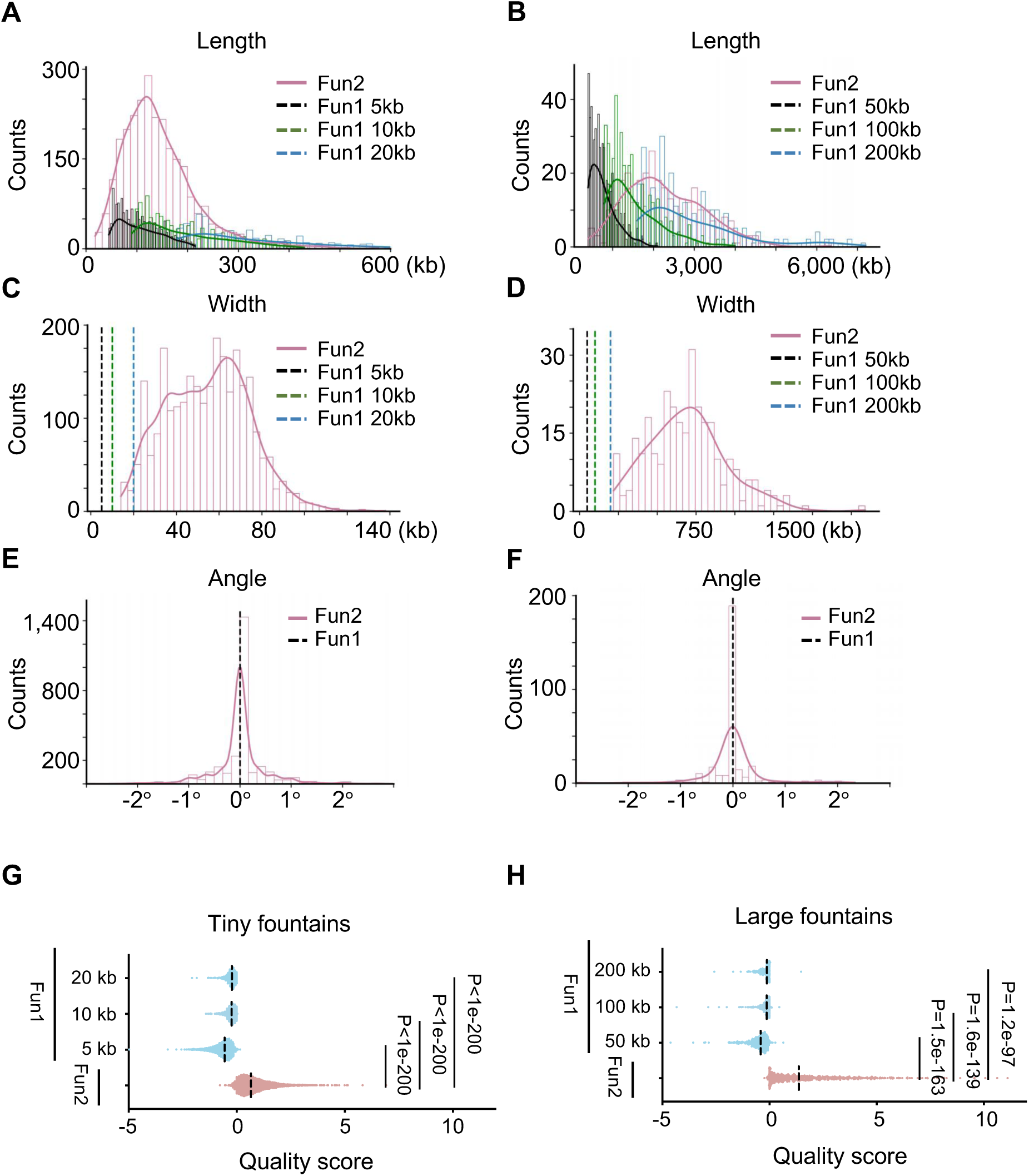
(A) The distribution of length for tiny-scale fountains identified by Fun1 and Fun2. The purple line with histogram indicates the results from Fun2, while the black, green, and blue dashed lines with histograms represent fountains identified at 5-kb, 10-kb, and 20-kb resolutions in Fun1, respectively. (B) The length distribution for large fountains, the purple line with histogram indicates the results from Fun2, while the black, green, and blue dashed lines with histograms represent fountains identified at 50-kb, 100-kb, and 200-kb resolutions in Fun1, respectively. (C) The width distribution for tiny-scale fountains, with the black, green, and blue dashed lines representing the fixed width of the sampling box in Fun1. (D) The width distribution for large fountains, with the black, green, and blue dashed lines representing the fixed width of the sampling box in Fun1. (E) The rotation angle distribution for tiny fountains. The black line indicates the fixed angle of the sampling box in Fun1. (F) The rotation angle distribution for large fountains. The black line indicates the fixed angle of the sampling box in Fun1. (G) The quality scores of identified tiny fountains in Fun1 (separately identified by data resolutions at 5-kb, 10-kb and 20-kb) and Fun2. One-sided Wilcoxon test. (H) The quality scores of identified tiny fountains in Fun1 (separately identified by data resolutions at 50-kb, 100-kb and 200-kb) and Fun2. One-sided Wilcoxon test.

**Supplementary Figure 8.**
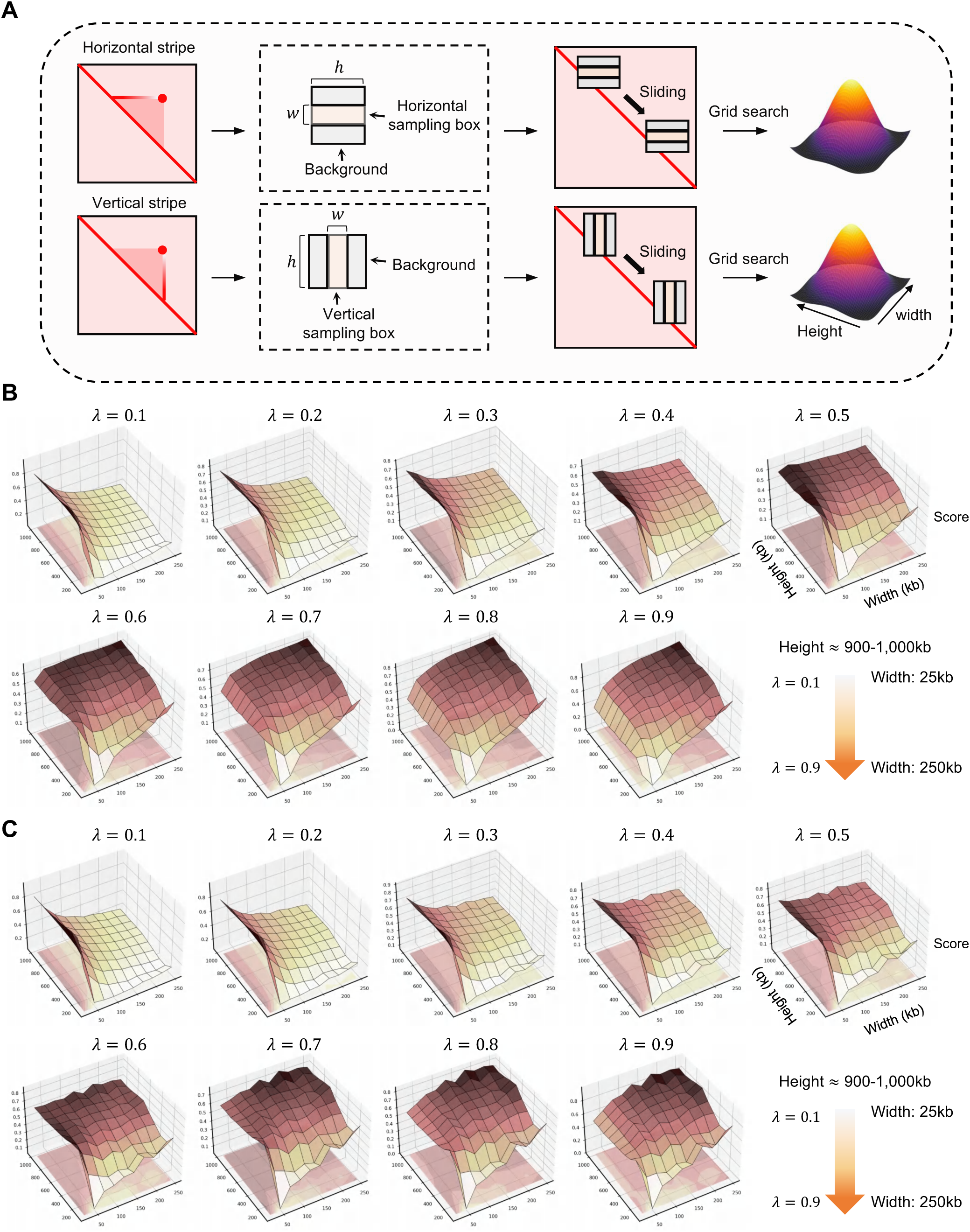
(A) Workflow of titration of summits for chromatin stripe detection. (B) Result of grid search for summits of horizontal chromatin stripes, with hyperparameter λ ranging from 0.1 to 0.9 is shown. For each λ, the length and width of the sampling box are adjusted within a defined range. (C) Result of grid search for summits of horizontal chromatin stripes, with hyperparameter λ ranging from 0.1 to 0.9 is shown. For each λ, the length and width of the sampling box are adjusted within a defined range.

**Supplementary Figure 9.**
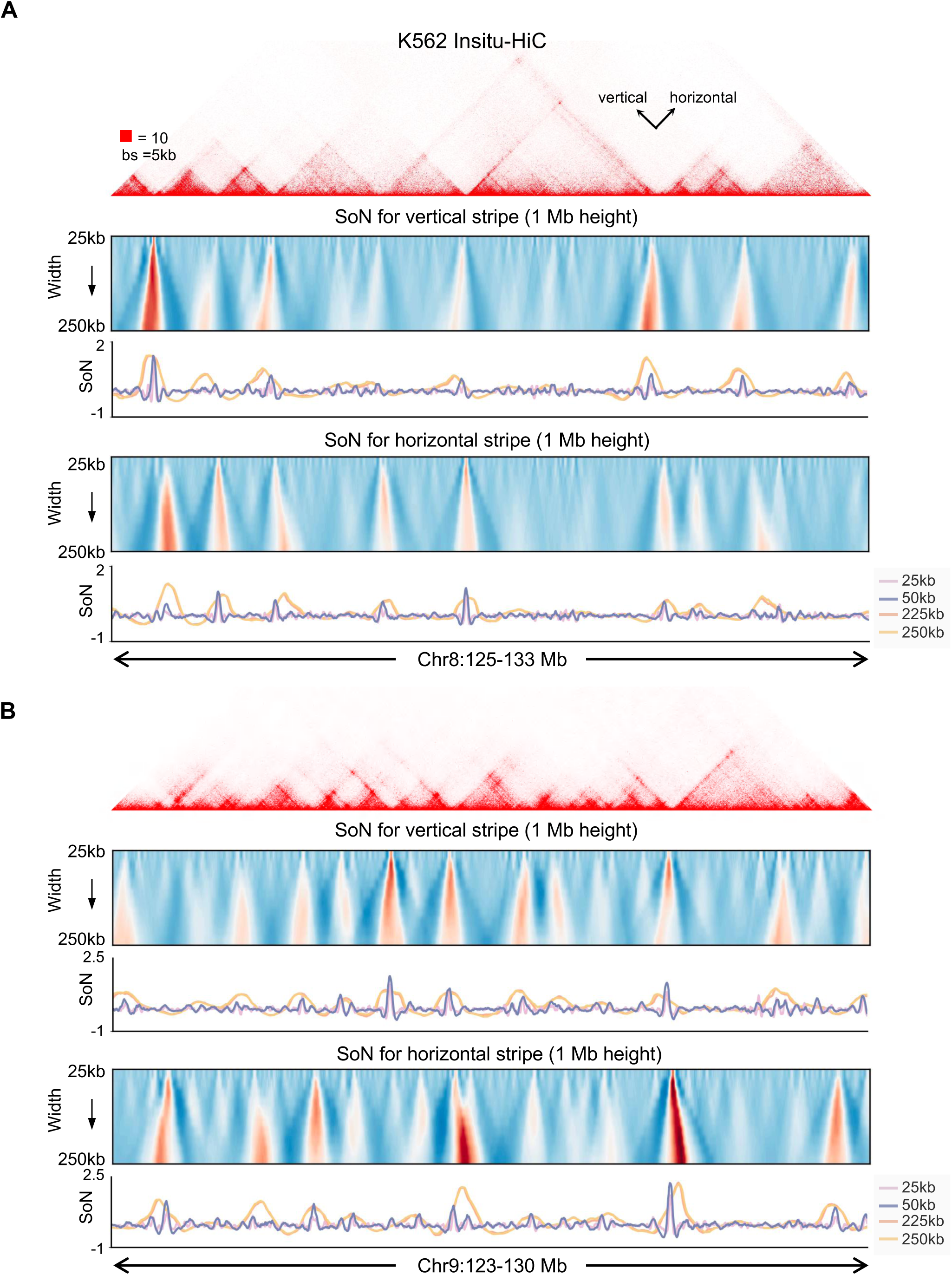
(A) An illustrative locus on Chr8 of K562 *in situ* Hi-C data with horizontal and vertical stripes. Heatmap showing the SoN signal calculated by sampling box with 1Mb height and width varying from 25kb to 250kb. Line plot showing stripes detection calculated by sampling box with 1Mb length and width of 25kb, 50kb, 225kb, 250kb. (B) An illustrative locus on Chr 9 of K562 *in situ* Hi-C data with horizontal and vertical stripes. Heatmap showing the SoN signal calculated by sampling box with 1 Mb height and width varying from 25 kb to 250 kb. Line plot showing stripes detection calculated by sampling box with 1 Mb length and width of 25 kb, 50 kb, 225 kb, 250 kb.

**Supplementary Figure 10.**
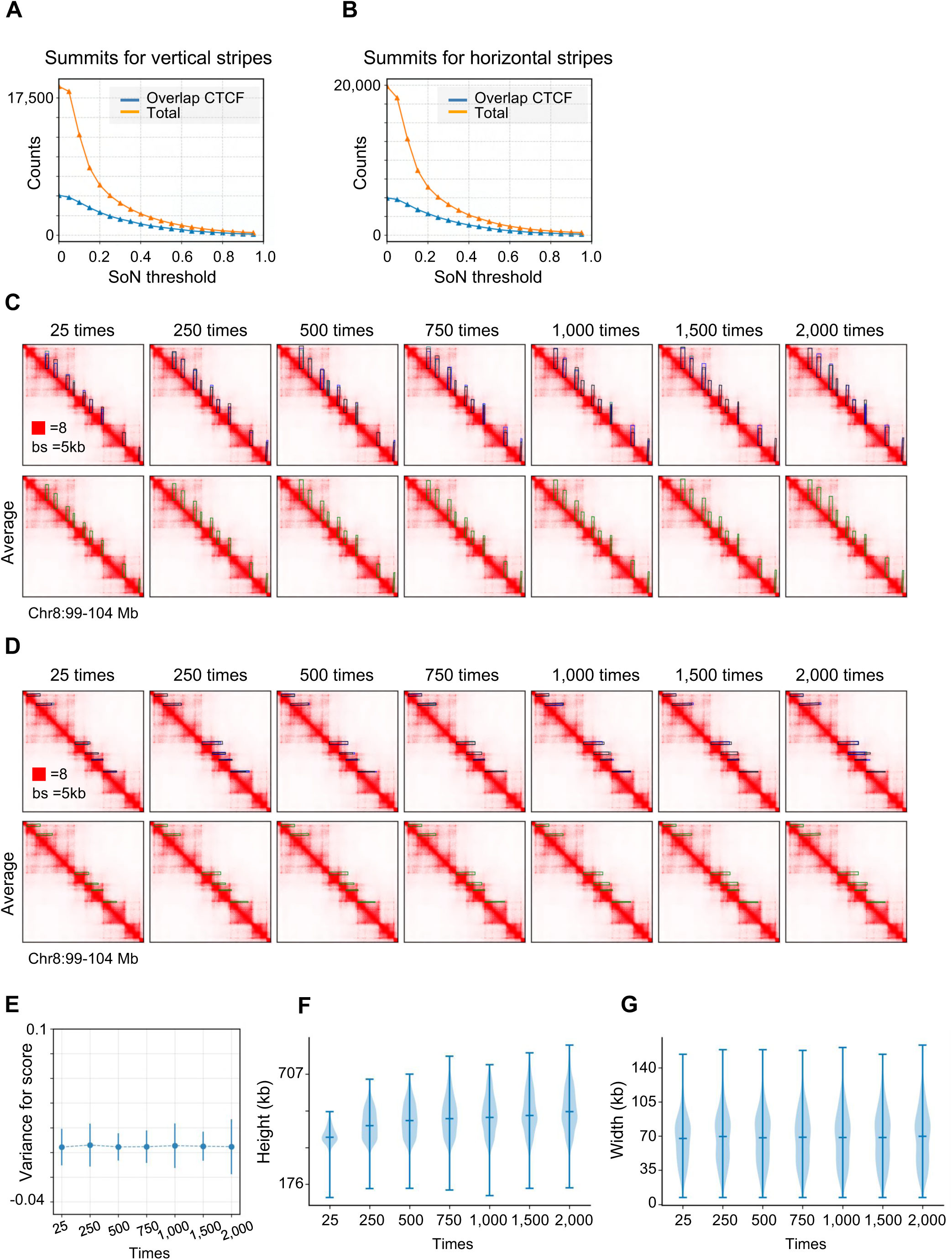
(A) Number of summits of vertical stripes at different SoN thresholds. The blue line indicates the number of summits that overlap CTCF enriched sites at varying SoN thresholds, while the orange line shows the total number of summits at each SoN threshold. (B) Same as (A) but for horizontal stripes. (C) An illustrative locus on Chr8 showing identified vertical chromatin stripes at training times varying from 25 times to 2,000 times. (D) Same as (C) but for horizontal stripes. (E) The variance in quality score of chromatin stripes on Chr8 across different training times, based on the average result. (F) The distribution of identified length of chromatin stripes at different training times. (G) The distribution of identified width of chromatin stripes at different training times.

